# Fine-scale cultural variation reinforces genetic structure in England

**DOI:** 10.1101/2022.09.23.509228

**Authors:** Yakov Pichkar, Nicole Creanza

## Abstract

Genes and languages both contain signatures of human history. Genetics and culture have each been shown to track population movements and demographic history. Complicating this picture, cultural traits may themselves influence the ways in which people interact with one another. For example, cultural differences can produce barriers to gene flow if they cause groups of people to differentiate themselves from one another. However, the degree of cultural difference necessary and the magnitude of these effects on gene flow remain unknown. In particular, language differences may limit population mixing, and we focus on whether subtle, dialect-level linguistic differences have influenced genetic population structure, likely by affecting mating preferences. Here, we analyze spatially dense linguistic and genetic data to examine whether the intensity of differences between and within dialects in England are associated with high genetic rates of change. We find that genetic variation and dialect markers have similar spatial distributions on a country-wide scale, and become less distinct as the scale of smaller administrative units such as counties. This covariation, combined with the absence of geographic barriers that could coordinate cultural and genetic differentiation, suggests that some dialect-level linguistic boundaries have influenced the genetic population structure in England.

## Introduction

For as long as evolution has been studied, researchers have worked to understand the relationship between biological traits and socially learned behaviors, particularly languages (1–4). There are striking similarities between genetics and culture, both of which spread and change as groups of people interact and move across the world (5, 6). When people move into a region, their genes and languages can begin to diverge from their previous population and mix with those of nearby populations (7). The resulting spatial patterns of language and genetics can mirror these demographic histories (8). Unlike vertically transmitted genetic traits, cultural traits are often inherited from people other than parents, so the transmission of cultural and biological traits can deviate from one another at the scale of individuals (9).

However, if similar individuals preferentially interact with one another—a tendency termed homophily—the population becomes structured, and this structure limits the transmission of traits (10–12). In animals, these assortative interactions can be genetically heritable due to selection against interspecific mating, but humans can also learn to bias their interactions in complex and subtle ways, resulting in homophily based on learned behaviors (13–15). When culture creates barriers to contact and interaction, it can contribute to covariation between genes and culture at relatively small scales. In this way, learned behaviors and preferences can act as cultural barriers to gene flow, thus altering the course of evolution.

Previous studies have investigated the distribution and the causes of this gene-culture covariation. In some cases, the evolution of cultural traits provides selective pressures that shape genetic variation. For example, the consumption of unfermented dairy products (16) or of high-starch diets (17) altered the selective pressures on genes related to metabolism. Other traits do not affect fitness directly, but can act as markers of identity that drive cultural homophily. One such trait is the specific set of languages or dialects a person speaks, since languages contain clues about the history of populations (6, 18–21) and have been found to co-vary with genetic variation on a worldwide scale (5, 6), though this pattern of covariation is not found everywhere (7, 22). Finer-scale analyses have found linguistic-genetic covariation in smaller regions as well, including the Caucasus (23, 24), the Levant (25), the Amazon (26), and Africa (27, 28). Other studies have directly compared the rates of linguistic and genetic change, as these are expected to be highest near geographic or behavioral barriers to the interaction of people (29). Such regions with populations differing in both languages and genetics have been found on the borders of countries in Europe (29), and even within countries (30).

These earlier studies observed evidence of gene-language covariation, but they have not been able to fully analyze its causes. Most of these studies compared people speaking different languages and did not use quantitative measures of language variation (29–31), though others have done this in their analyses (6, 20, 32), and recent datasets have attempted to digitize cultural information and linguistic relationships on a global scale (33, 34). The limits of previous studies were due, in part, to the limited availability of quantitative data describing cultural variation and the low spatial resolution of available genetic data. As a result, previous work in this field has only compared genes and language at the scale of ethnic groups and large regions, where the effects of population movement and cultural homophily cannot be disentangled. As such, the impact and spatial extent of these forces has not been thoroughly studied, especially at the smaller, intra-population scales of interaction that are predominant in the lives of most people.

To elucidate the role of homophily in contributing to gene-language covariation, we present a joint analysis of genes and language in England. We use fine-scale linguistic and genetic data to measure the intensity, scale, and location of covariation. To compare genetics and language, we analyzed two separate sets of data: the Survey of English Dialects (SED) (35) and the People of the British Isles (PoBI) project (36). Each dataset has a high geographic density, has been used previously to study the history of the region, and independently captures spatially meaningful linguistic and genetic variation (32, 36, 37). Leslie et al. (36) used the PoBI dataset to generate genetic similarities and clustered individuals into groups based on these similarities; these clusters retain evidence of population movements within the British Isles and from different parts of mainland Europe. These recently gathered and digitized data allow us to use quantitative methods to investigate how culture has shaped the genetic structure of England over the past thousand years.

By using these genetic and language data in concert, we measure the relationship between genes and culture that formed over millennia in Great Britain. Comparing the rates of linguistic and genetic change throughout England, we find similarities between their spatial distribution at all scales of analysis, suggesting that subtle cultural boundaries could reinforce genetic population structure.

## Results

We added to a digitized database of features (32) from the Survey of English Dialects (SED) to quantify linguistic variation by measuring the total rates of change across all of these phonetic features in a region (Fig. 1). Regions in which nearby locales have differences across many features suggest the presence of a border between dialects, and also, of a difference in the underlying cultural identities of people living there. We observed patterns of linguistic variation similar to those found in other analyses of the SED (35, 38). Our linguistic clustering identified the geographic distribution of features associated with dialectics in England, and the areas where these clusters meet also generally show high rates of language change (Fig 1.). We found high rates of language change near the boundaries of several dialect areas, namely near the Scottish Border, the southern Welsh Marches, and between the West Country and the rest of South England.

**Fig. 1.**
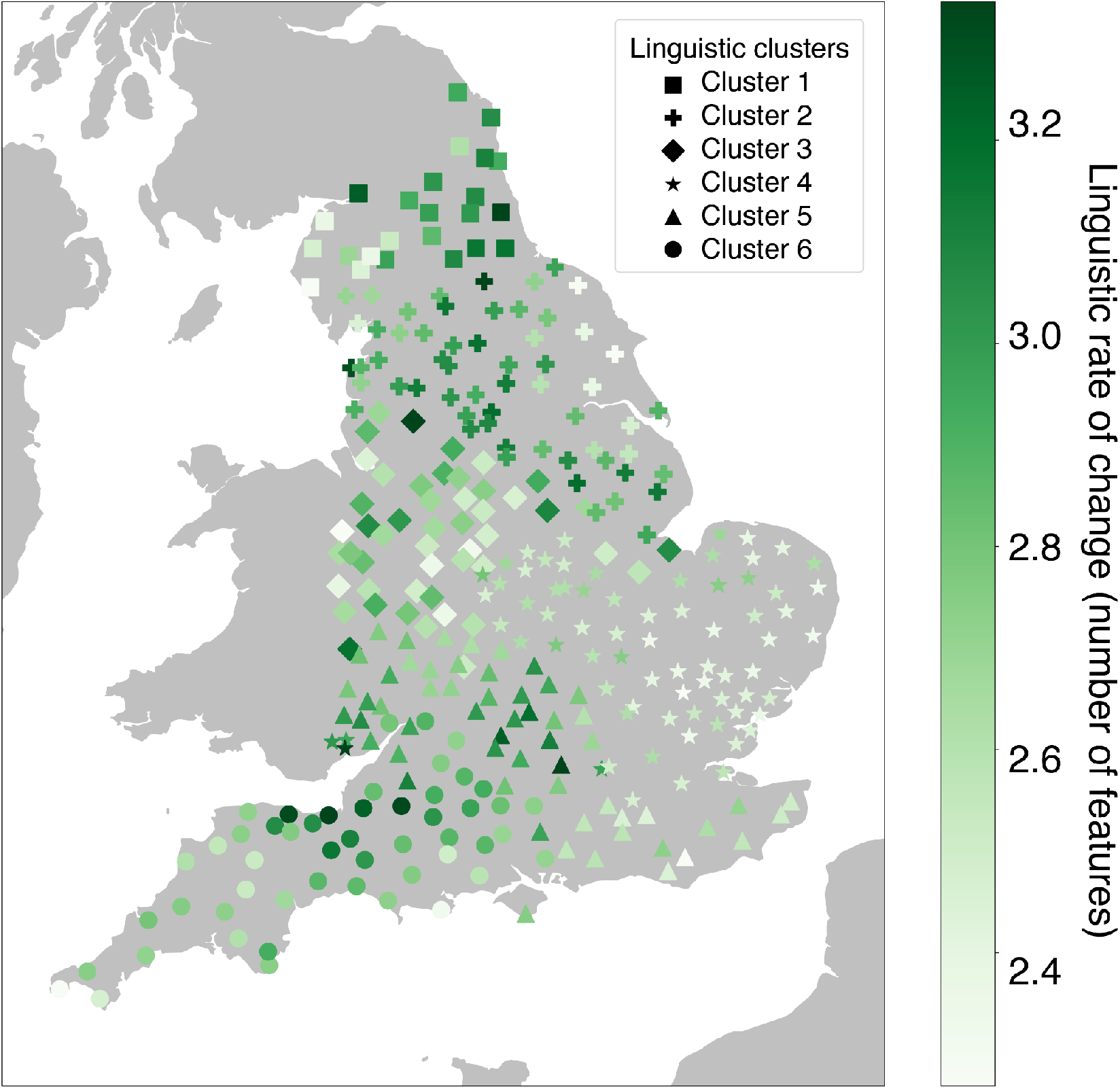
Linguistic rates of change in England. For each location, we estimated the linguistic rates of change by finding the number of linguistic features that differ between that location and those within 100 km of it, and finding their average, weighted by the inverse of the spatial distances to those locations. We performed hierarchical clustering on the linguistic data (*K*=6), and different symbols show which samples were members of each cluster.

We next used genomic data from the People of the British Isles (PoBI) study and fineSTRUCTURE to group 1,667 individuals from England into six geographically distinct hierarchical clusters. These were very similar to the clusters found in England by Leslie et al. (36), but they are not identical due to the stochastic nature of fineSTRUCTURE. The largest of these clusters was dominant in South England and the Midlands, containing nearly half of all sampled individuals. Other clusters were found near the Welsh border, the Scottish border, Northern England (centered on Yorkshire), Somerset, and Cornwall (Fig. 2).

**Fig. 2.**
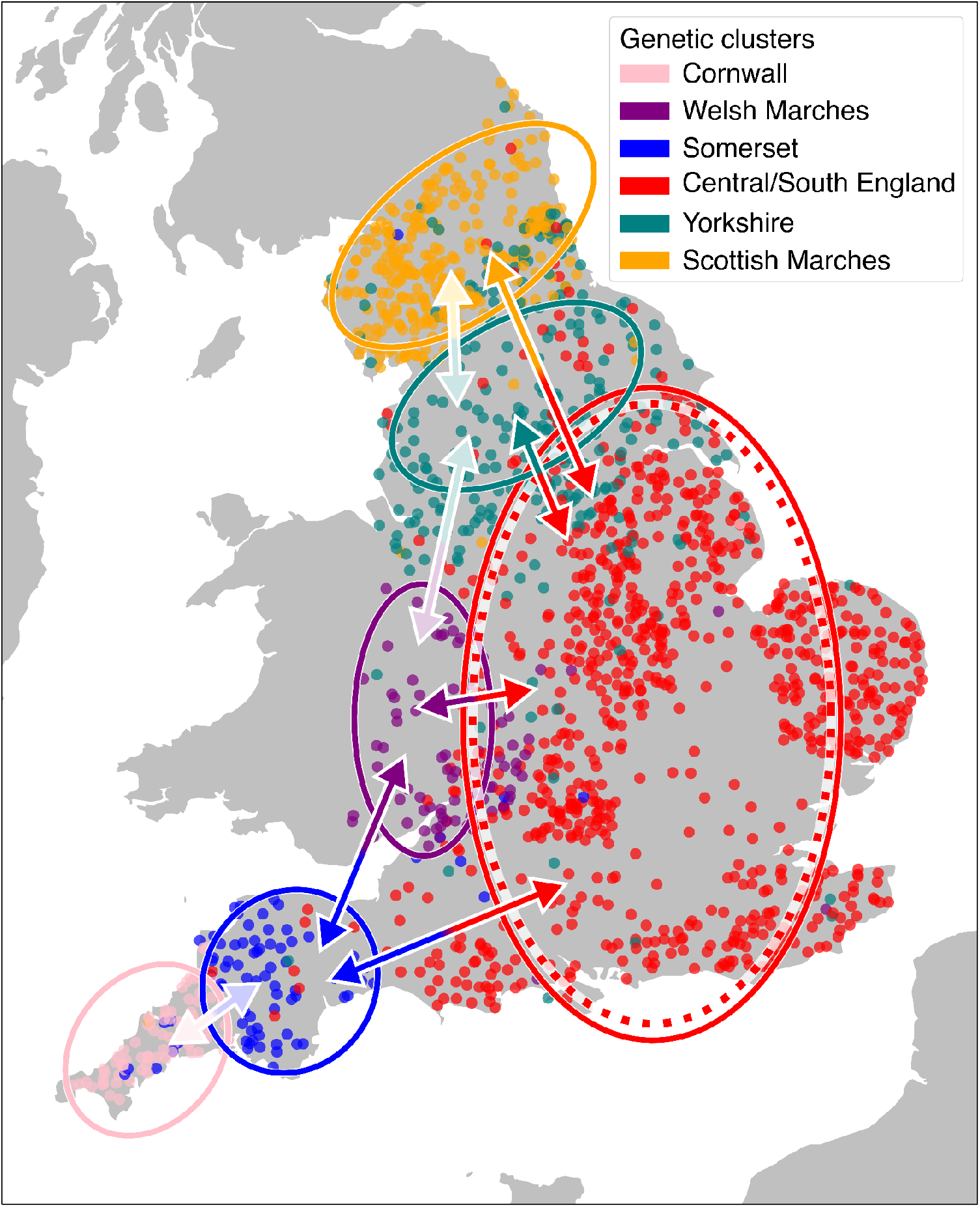
Relationship between genetic and linguistic rates of change. For between-cluster boundaries (double-sided arrows), a significant gene-language relationship is shown by a dark arrow and a non-significant relationship is shown by a light arrow. Specifically, we measured whether regions with high rates of genetic change—particularly cluster boundaries—had higher rates of linguistic change, corresponding to dialectal and other cultural boundaries. We draw a dotted ellipse in the cluster that represents much of the Midlands and South England, where there is a significant correlation between the distributions of genes (of individuals in that cluster) and language (in the same region). No other clusters have this relationship (see Table 1). In this figure, the hierarchical clustering was set to *K*=6.

**Table 1.** Genetic-linguistic covariation between genetic clusters. For each set of adjacent clusters, we measured whether their cluster boundary was found in regions with high linguistic rates of change using Mann-Whitney *U* tests. We also include the number of locations used for the tests. Bold values indicate those significant after Bonferroni correction.

| Cluster 1 | Cluster 2 | $N_1, N_2$ | $U$-value | $p$-value |
| --- | --- | --- | --- | --- |
| Somerset | Cornwall | 24, 7 | 131.0 | 0.99 |
| Somerset | Welsh Marches | 62, 15 | 185.0 | **$1.6 \times 10^{-4}$** |
| Yorkshire | Scottish Marches | 82, 48 | 1531.0 | $1.76 \times 10^{-2}$ |
| South/Central England | Scottish Marches | 213, 66 | 4724.0 | **$3 \times 10^{-5}$** |
| South/Central England | Yorkshire | 208, 64 | 4581.0 | **$8 \times 10^{-5}$** |
| Yorkshire | Welsh Marches | 126, 29 | 2471.0 | 0.999 |
| South/Central England | Welsh Marches | 221, 62 | 3356.0 | **$< 1 \times 10^{-5}$** |
| South/Central England | Somerset | 240, 25 | 1368.0 | **$< 1 \times 10^{-5}$** |

We used Procrustes analysis to measure the large-scale associations among geography, genetic variation, and linguistic variation in England. Based on these analyses, we found that dialect-level linguistic variation has a strong geographic signal (Procrustes correlation of 0.888, *p* < 10^−6^) and that it contains both north-south and east-west clines, though these are not smooth throughout England (Figs. 3B & S3C). Genetic variation is also correlated to geography (Procrustes correlation of 0.683, *p* < 10^−6^) and the genetic clusters form distinct groups in PC space (Fig. 3A). Genetics and linguistics covary, though to a lesser degree (Procrustes correlation of 0.531, *p* < 10^−6^). Most genetic clusters do not align with any specific linguistic cluster, but many have a linguistic cluster with which they are predominantly associated (Fig. 3A). For example, linguistic cluster 6 is dominant in Somerset and Cornwall (pink and blue circles) and linguistic cluster 1 is most common in the Scottish Marches (orange squares).

**Fig. 3.**
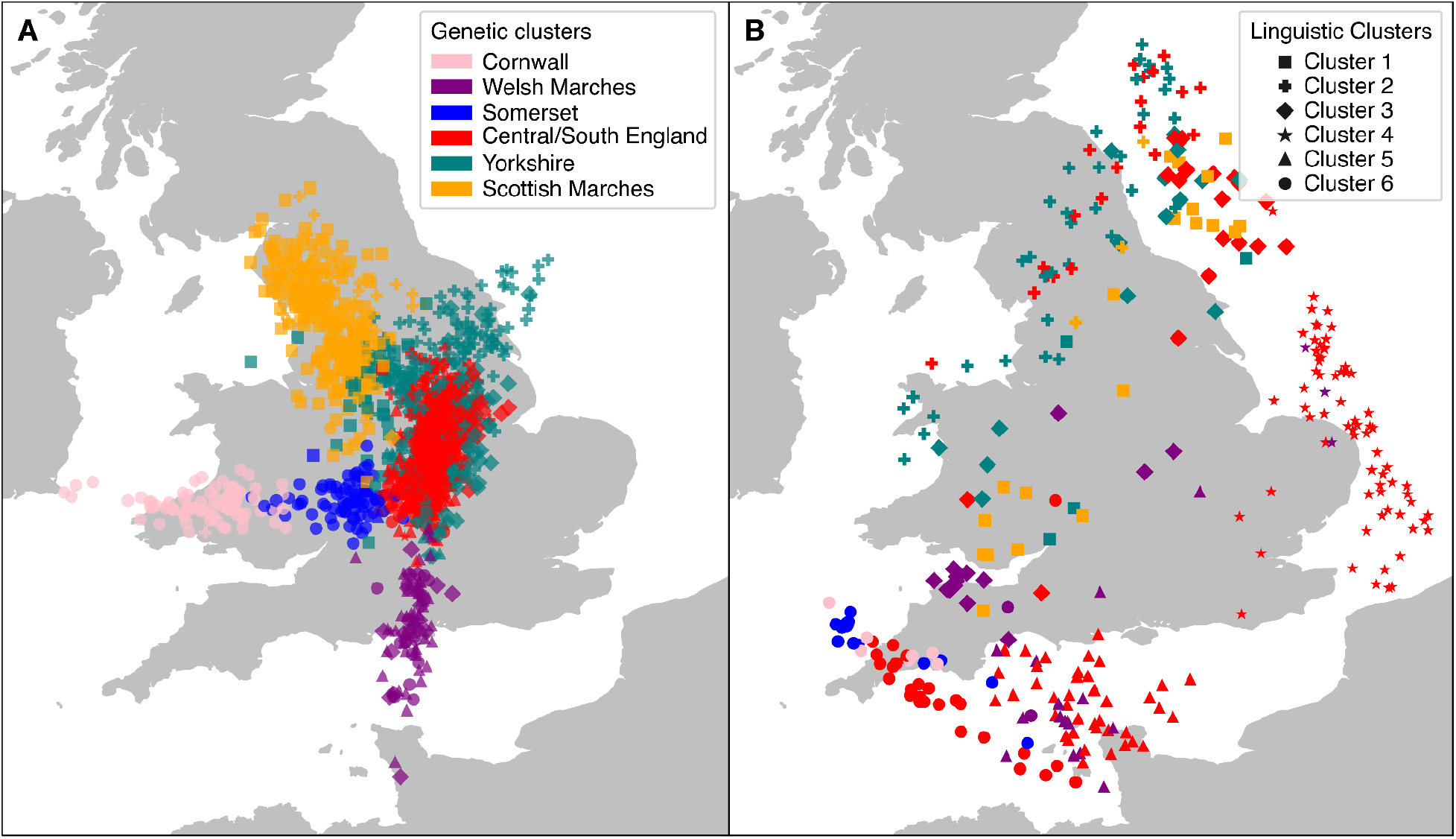
Procrustes transformations of linguistic and genetic data onto geography. We aligned geography to the first three principal components (PCs) of linguistic (A) and genetic (B) data for Procrustes analysis. The linguistic data is color-coded using approximate genetic clusters, and shapes represent linguistic clustering. The gene-geography Procrustes is color-coded using genetic clusters, and shapes represent approximate linguistic clusters. Both linguistic and genetic data are associated with geography (Procrustes correlations of 0.888 and 0.683, *p* < 10^−6^), but neither form of variation smoothly recapitulates the geographic distributions of sampling locations. Genetics and linguistics are correlated with one another, though this correlation is weaker than between either and geography (Procrustes correlation of 0.531, *p* < 10^−6^). In Fig. S6, we show these plots color-coded by geographic location instead of genetic cluster membership.

We find that linguistic rates of change are significantly greater at the boundaries between genetic clusters than at the cores of these clusters (Fig. 2 and Table 1). Across many regions of England, we identified that sharp differences in linguistics colocalized with genetic cluster boundaries. Regions with long-standing cultural differences—near the Welsh and Scottish borders—have co-occurring high rates of genetic and linguistic change (Fig. 2). Within the interior of England, in areas that have been historically English (those farther from borders with other countries), we also find a gene-language covariation. Between South/East England and both Yorkshire and the West Country, there are associations between genetic cluster boundaries and linguistic dialect boundaries. Despite Cornwall’s distinct ethnic history, dialect, and genetics, we do not find an association between genes and languages at its border with Somerset. However, the region’s unique dialect is most prominent in the westernmost end of the county, whereas our genetic data has at most a county-scale resolution (Figs. 1 and 2).

In addition to measuring how linguistic rates were associated with genetic cluster boundaries, we measured whether genetic variation within relatively well-mixed clusters co-varies with language. For each of our six clusters, we compared the rates of linguistic and genetic change within the boundaries of each cluster (Fig. 2, Table 2). Within the cluster containing most individuals in the Midlands to East England, there is a strong relationship between the spatial distribution of genetic and linguistic variation. Here—and in no other cluster—we found that genetic and linguistic rates of change are strongly correlated. The genetic variation here was not captured in our hierarchical clustering, as increasing the number of clusters beyond 6 could not divide the individuals into geographically distinct groups (Fig. S5). In Central and Southern England, this gene-language pattern is found at all scales of the hierarchical clustering (red cluster in Figs. S1-S5, Table S3), pointing to spatially heterogeneous genetic variation that is not inherent in the clustering.

**Table 2.**
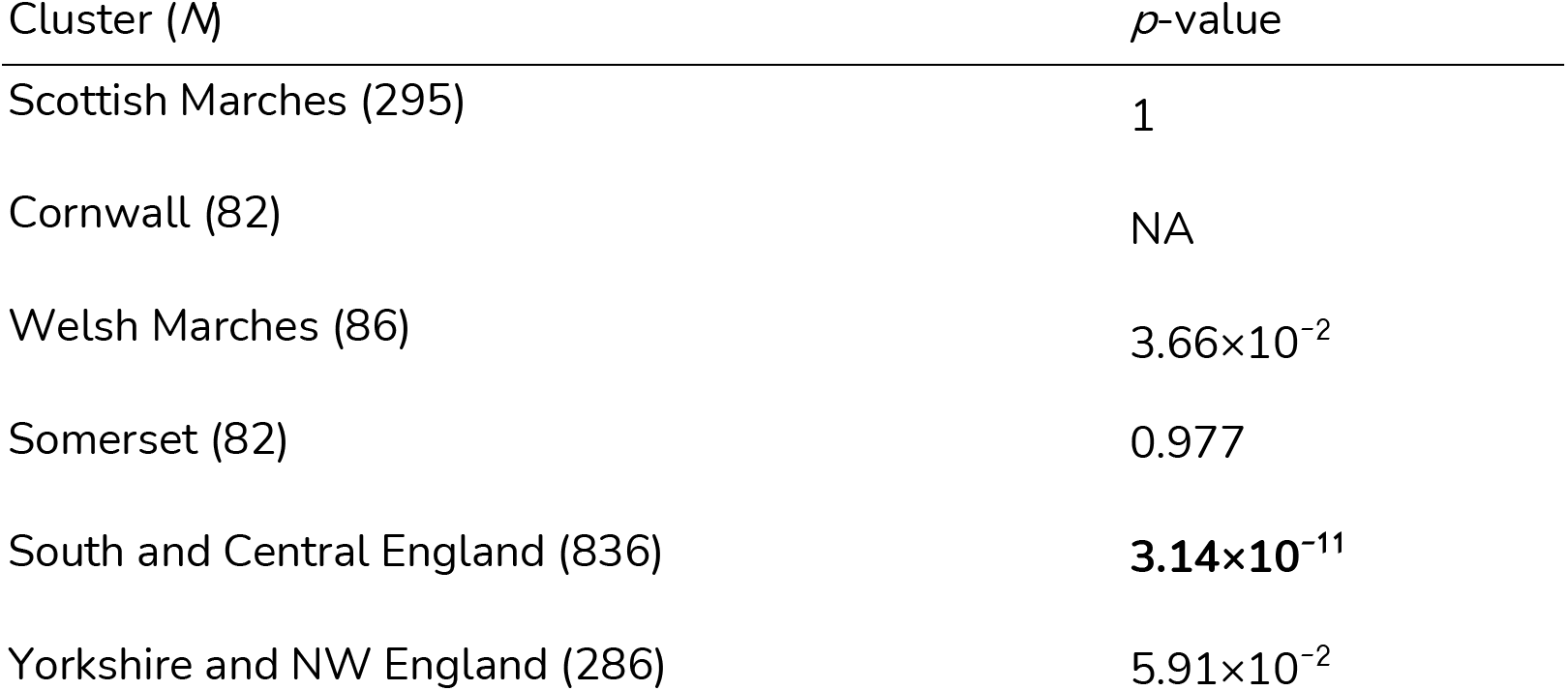
Genetic-linguistic covariation within clusters. For each cluster identified at *K* =6 hierarchical clusters, we compared genetic and linguistic rates of change using linear regression. Only a single cluster had significantly correlated genetic and linguistic rates of change. All of the individuals in the Cornwall genetic cluster were in a single county, so we could not compare their rates of genetic change. Bold values are those significant after Bonferroni correction. We include the number of individuals in each cluster (*N*).

## Discussion

Our analyses of genetic and linguistic data identify geographic patterns also found in previous studies, and they show gene-culture interactions at the dialect level that were only previously observed at the language level. We measured the spatial distribution for rates of language change throughout England, including near various cultural boundaries (35, 38). We found high rates of language change at the borders of England and at the edges of other previously identified dialects within England (Fig. 1), suggesting that our measure is a good proxy for cultural variation. Our genetic analysis also identified spatial patterns similar to those found previously. These genetic patterns reflect waves of migration to Great Britain and of later movements of people (39). We found six geographically distinct genetic clusters ranging in size from 82 to 836 individuals (Figs. 2 and 3); these clusters spanned regions as small as a single county (Cornwall) to large enough to encompass most of the Midlands and South England. Attempting to cluster this genetic data into more than six clusters with fineSTRUCTURE did not reveal more geographic population structure (Fig. S5). Both the genetic and language data covaried with geography (Figs. 3 and S6), pointing to their general spatial patterns and potential for informing our analyses.

We find that linguistic and genetic variation parallel one another throughout much of England. The boundaries of genetic clusters almost always appear in regions with high rates of cultural change. This correspondence between genetic boundaries—which are associated with both historical migrations and modern distributions of the peoples of Great Britain—and linguistic boundaries—associated with dialect and other cultural differences—demonstrate that language and genetics have similar geographic patterns in England. These gene-language patterns at the English-Welsh and the English-Scottish borders support previous studies that identify the covariation of culture and genetics near country borders and ethnic boundaries (23, 25, 27–30). In addition to regions with prominent cultural differences at the borders of England, we find that this covariation extends to less dramatic distinctions, such those between South East and South West England or between North and South England. These genetic clusters are not only geographically localized, but also contain variation associated with various admixture events from mainland Europe, including the arrival of the Celts, Anglo-Saxons, Danes, and Normans (36). Our results suggest that cultural distinctions, such as dialect differences, have influenced how people move and behave by maintaining genetic boundaries; however, the results cannot fully distinguish between the influence of cultural homophily and that of other demographic processes, both of which may have contributed to the existing patterns of genes and language.

To find evidence of cultural barriers to gene flow, we also analyzed the genetic variation existing within genetic clusters, rather than considering only boundaries between clusters. We tested whether regions with higher rates of language change within a genetic cluster corresponded to higher rates of genetic change within that cluster. We find that, for smaller clusters, genetic differences do not appear to covary with rates of language change. However, we find this relationship for the largest cluster, which encompasses much of the Midlands and South England. Despite this cluster appearing relatively well-mixed, in the sense that it cannot be divided into more geographically meaningful clusters, higher rates of genetic change coincide with dialect boundaries. These data suggest that dialect boundaries—or related cultural differences—contribute to genetic differentiation in that region. The lack of major natural barriers to human movement in Central and South England imply that the population should be relatively homogeneous; however, the varied history of the Midlands and South England (36) and existing covariation between barriers to gene flow and dialect boundaries strongly suggest that cultural variation has been responsible for shaping the genetics of the region.

The presence of gene-culture covariation and the longevity of these genetic clusters—which contain characteristics of movements in the past—point to the role of culture in maintaining genetic variation. For genetic variation within clusters to covary with culture, either a third force (such as geography) limited their diffusion, or cultural differences contributed to the current genetic population structure. Given the many roads and navigable rivers in England (40), it is likely that cultural homophily has acted as a major barrier to gene flow.

Our analysis of the interplay between genetics and culture has greater spatial resolution and more quantitative data than most previous studies (24, 28–30), but we are still limited by the resolution of our data and our use of separate sets of data. Our genetic data was restricted to spatial resolutions as fine as counties, which prevented the comparison of genetics to intra-county linguistic variation. Since we used separate genetic and linguistic datasets, we could only comment on processes that affect both of these over many generations. Ideally, we would more directly measure assortative mating based on cultural homophily, which could be possible with cultural and lifetime decision-making data from a set of genotyped individuals. Although we cannot directly measure homophily, our results are consistent with previous work that identified widespread endogamy and geographically restricted marriages in England (41). Such homophily likely had profound effects on population structure (42–44), and we suggest that cultural distinctions, through homophily, have shaped the genetics of England. Similar processes have influenced the ways in which families form and ideas spread, across the world and throughout time. The small scale at which we have identified cultural homophily suggests that cultural divergence is a major force that structures human populations, and this should be taken into account when designing genetic studies.

## Materials and Methods

### Processing linguistic data

Linguistic data was collected by the Survey of English Dialects (SED) (35), which gathered word choice and pronunciation in England, the Isle of Man, and parts of Wales. The SED, conducted in the 1950’s, prioritized people who were elderly, from rural areas, and who had agricultural backgrounds. This survey took place across 313 locales, of which we excluded two on the Isle of Man, for which we have no genetic data.

From the SED, we digitized 225 binary elements that describe whether a particular sound is present in each word, expanding our previous digitized SED data of 45 elements (Table S1) (32). For example, an element may be whether “hammer” is pronounced with or without an initial [h] sound.

We took these 225 binary elements and combined those that were relevant into 55 categories representing the frequencies of specific linguistic features, such as the presence of syllable-initial [h] (the mean occurrence of syllable-initial [h] in hammer, halter, harvest, etc., Table S1). If a linguistic feature contained multiple binary elements, each of the 311 localities was assigned a frequency of occurrence for that feature; if a linguistic feature was represented by only a single element, the data were binary for that category.

In addition, we divided the linguistic data into clusters by finding the pairwise standardized Euclidean distance between locations and using the Ward criterion to create groups.

### Processing genetic data

We used genetic data collected for the People of the British Isles (PoBI) project, which included the genotyping of people from the British Isles. These people were mostly from rural backgrounds and all had grandparents born within 80 km of one another; these grandparents had a mean birth year of 1885 (s.d.=18yrs.). The genetic data consists of over 500,000 single nucleotide polymorphisms (SNPs) for each of 2,039 people, after quality control. Each of these individuals was placed into one of 36 locations in Great Britain and Northern Ireland to protect their anonymity, and 29 of these are in England. Previous work by Leslie et al. found that these individuals could be placed into 17 geographically cohesive clusters based on their genetics.

We replicated the quality control, phasing, and clustering procedure from Leslie et al. Since our linguistic data was limited to England, we used a subset of 1,667 individuals by keeping those whose grandparents were born in England. This differs from the data used by Leslie et al., as they included all 2,039 individuals with grandparents from England, Wales, Scotland, and Northern Ireland (36).

After these filters, we used fineSTRUCTURE to hierarchically cluster these individuals into geographically cohesive groups based on genetics. Beginning with two clusters, we increased the number of clusters until we found six such clusters; past that point, new clusters lost geographic cohesiveness. These six clusters correspond to the six English clusters found by Leslie et al. within England.

### Gene-language processing

Since the genetic and linguistic datasets include information from different individuals, we conducted our analyses based on the geographic sampling locations of the individuals in each dataset. We assigned each of the 311 locales with linguistic data to one of the 29 county-level regions with genetic data. Each locale of these 311 locations was treated as including individuals with genetic gata from that region.

We found the Euclidean distance between the linguistic features for all pairs of locations within 100 km of one another. We then found the average rate of linguistic change at that location by averaging the linguistic distances to that location, weighed by the inverse of spatial distances to that location.

To find genetic distances, we used the output produced by fineSTRUCTURE, termed a genetic copying matrix, which represents the total recombination map distance of haplotypes donated from one individual to another, according to fineSTRUCTURE’s model (45). Distances between copying vectors have been used previously to find the genetic distances between groups of individuals (36). We used this method to calculate distances between each pair for individuals, which we then used to find the mean distance between clusters and between regions with genetic data. We did this for every pair of adjacent genetic clusters from fineSTRUCTURE, and within every cluster found in more than one location. For every set of clusters being compared, for each pair of locales within 100 km of one another, we found the mean genetic distance between individuals from different clusters (or within the same cluster for within-cluster comparisons) and from the genetic regions associated with those locales. We found rates of genetic change at each location using genetic distances, identically to rates of linguistic change.

We repeated the steps to calculate genetic rates of change 2000 times, randomizing the order of the genetic relatedness matrix each time. Using these rates with randomized data, we computed the true rates’ percentile to find a standardized rate of genetic change, making them comparable and to decreasing sampling biases.

To compare the genetic and linguistic rates of change, we used two methods. First, to evaluate this relationship within clusters, we measured whether a positive relationship existed via linear regression of the linguistic and genetic rates of change. We calculated these regressions for each genetic cluster which contained individuals from multiple counties (i.e. excluding the cluster found only in Cornwall). Second, to measure the overlap of genetic cluster boundaries and dialect boundaries, we used the Mann-Whitney U test to assess whether linguistic rates of change were greater at cluster boundaries than they were in other regions in which the clusters appeared.

### Procrustes analysis

To compare the distributions of linguistic and genetic variation, we found the Procrustes transformations that best aligned their variation to geography. We calculated the first three principal components (PCs) for both the genetic data (using the genetic copying matrix) and linguistic data (using the linguistic feature frequencies). For each data type, we could then compare their PCs to their geographic locations in Britain (converted to Cartesian coordinates) and find their Procrustes correlations. Since we had 311 samples in the linguistic database, we
 repeatedly sampled 311 of the 1,667 genotyped individuals before calculating genetic principal components 500 times to compare the gene-geography and language-geography Procrustes with one another. We also used this resampling method when comparing genetic and linguistic variation. To measure significance of each Procrustes analysis, we permuted the order of the data 10^6^ times, and we calculated an empirical *p*-value as the fraction of permuted samples that had greater Procrustes correlations than the original data.

## Acknowledgments

We thank Walter Bodmer, Sohini Ramachandran, Marc Feldman, and members of the Creanza lab for helpful discussions and Andre Sherriah, Hubert Devonish, and Sydnie Smith for help with data coding. Funding was provided by Vanderbilt University, Vanderbilt’s Evolutionary Studies Institute, and the John Templeton Foundation.

## Notes

### Competing Interest Statement

The authors have declared no competing interest.

### Summary of Updates

New analyses and figures added.

## References

1. C. Darwin, The Descent of man (1871).

2. L. L. Cavalli-Sforza, M. W. Feldman, Cultural transmission and evolution: a quantitative approach. Monogr. Popul. Biol. 16, 1–388 (1981).

3. C. J. Lumsden, E. O. Wilson, GENES, MIND, AND IDEOLOGY. The Sciences 21, 6–8 (1981).

4. R. Boyd, P. J. Richerson, Culture and the Evolutionary Process (University of Chicago Press, 1985).

5. L. L. Cavalli-Sforza, E. Minch, J. L. Mountain, Coevolution of genes and languages revisited. Proc. Natl. Acad. Sci. U. S. A. 89, 5620–5624 (1992).

6. N. Creanza, et al., A comparison of worldwide phonemic and genetic variation in human populations. Proc. Natl. Acad. Sci. U. S. A. 112, 1265–1272 (2015).

7. N. Creanza, M. W. Feldman, Worldwide genetic and cultural change in human evolution. Curr. Opin. Genet. Dev. 41, 85–92 (2016).

8. M. J. Hamilton, B. Buchanan, Spatial gradients in Clovis-age radiocarbon dates across North America suggest rapid colonization from the north. Proc. Natl. Acad. Sci. U. S. A. 104, 15625–15630 (2007).

9. C. R. Guglielmino, C. Viganotti, B. Hewlett, L. L. Cavalli-Sforza, Cultural variation in Africa: role of mechanisms of transmission and adaptation. Proc. Natl. Acad. Sci. U. S. A. 92, 7585–7589 (1995).

10. N. Creanza, M. W. Feldman, Complexity in models of cultural niche construction with selection and homophily. Proc. Natl. Acad. Sci. U. S. A. 111 Suppl 3, 10830–10837 (2014).

11. E. Katsnelson, A. Lotem, M. W. Feldman, Assortative social learning and its implications for human (and animal?) societies. Evolution 68, 1894–1906 (2014).

12. F. Fu, M. A. Nowak, N. A. Christakis, J. H. Fowler, The evolution of homophily. Sci. Rep. 2, 845 (2012).

13. I. Eshel, L. L. Cavalli-Sforza, Assortment of encounters and evolution of cooperativeness. Proc. Natl. Acad. Sci. U. S. A. 79, 1331–1335 (1982).

14. A. Abdellaoui, et al., Association between autozygosity and major depression: stratification due to religious assortment. Behav. Genet. 43, 455–467 (2013).

15. A. Tenesa, K. Rawlik, P. Navarro, O. Canela-Xandri, Genetic determination of height-mediated mate choice. Genome Biol. 16(2015).

16. A. Beja-Pereira, et al., Gene-culture coevolution between cattle milk protein genes and human lactase genes. Nat. Genet. 35, 311–313 (2003).

17. R. C. Iskow, O. Gokcumen, C. Lee, Exploring the role of copy number variants in human adaptation. Trends Genet. 28, 245–257 (2012).

18. C. Barbieri, A. Butthof, K. Bostoen, B. Pakendorf, Genetic perspectives on the origin of clicks in Bantu languages from southwestern Zambia. Eur. J. Hum. Genet. 21, 430–436 (2013).

19. C. Barbieri, et al., Between Andes and Amazon: the genetic profile of the Arawak-speaking Yanesha. Am. J. Phys. Anthropol. 155, 600–609 (2014).

20. P. Verdu, E. M. Jewett, T. J. Pemberton, N. A. Rosenberg, M. Baptista, Parallel Trajectories of Genetic and Linguistic Admixture in a Genetically Admixed Creole Population. Current Biology 27, 2529–2535.e3 (2017).

21. R. D. Gray, A. J. Drummond, S. J. Greenhill, Language phylogenies reveal expansion pulses and pauses in Pacific settlement. Science 323, 479–483 (2009).

22. K. L. Hunley, G. S. Cabana, D. A. Merriwether, J. C. Long, A formal test of linguistic and genetic coevolution in native Central and South America. Am. J. Phys. Anthropol. 132, 622–631 (2007).

23. T. M. Karafet, et al., Coevolution of genes and languages and high levels of population structure among the highland populations of Daghestan. J. Hum. Genet. 61, 181–191 (2016).

24. O. Balanovsky, et al., Parallel evolution of genes and languages in the Caucasus region. Mol. Biol. Evol. 28, 2905–2920 (2011).

25. M. Haber, et al., Genome-wide diversity in the levant reveals recent structuring by culture. PLoS Genet. 9, e1003316 (2013).

26. T. Di Corcia, et al., East of the Andes: The genetic profile of the Peruvian Amazon populations. Am. J. Phys. Anthropol. 163, 328–338 (2017).

27. E. G. Atkinson, et al., Genetic structure correlates with ethnolinguistic diversity in eastern and southern Africa. Am. J. Hum. Genet. 109, 1667–1679 (2022).

28. D. Sengupta, et al., Genetic substructure and complex demographic history of South African Bantu speakers. Nat. Commun. 12, 2080 (2021).

29. G. Barbujani, R. R. Sokal, Zones of sharp genetic change in Europe are also linguistic boundaries. Proc. Natl. Acad. Sci. U. S. A. 87, 1816–1819 (1990).

30. G. Barbujani, R. R. Sokal, Genetic population structure of Italy. II. Physical and cultural barriers to gene flow. Am. J. Hum. Genet. 48, 398–411 (1991).

31. T. M. Karafet, et al., Coevolution of genes and languages and high levels of population structure among the highland populations of Daghestan. J. Hum. Genet. 61, 181–191 (2016).

32. A. C. Sherriah, H. Devonish, E. A. C. Thomas, N. Creanza, Using features of a Creole language to reconstruct population history and cultural evolution: tracing the English origins of Sranan. Philos. Trans. R. Soc. Lond. B Biol. Sci. 373 (2018).

33. S. Passmore, et al., Global relationships between musical, linguistic, and genetic diversity https://doi.org/10.31234/osf.io/mdrsn.

34. K. R. Kirby, et al., D-PLACE: A Global Database of Cultural, Linguistic and Environmental Diversity. PLoS One 11, e0158391 (2016).

35. H. Orton, E. Dieth, Survey of English Dialects (1963).

36. S. Leslie, et al., The fine-scale genetic structure of the British population. Nature 519, 309–314 (2015).

37. C. Upton, J. D. A. Widdowson, An Atlas of English Dialects: Region and Dialect (Routledge, 2013).

38. W. Viereck, Dialectal speech areas in England: Orton’s phonetic and grammatical evidence. J. Eng. Linguist. 19, 240–257 (1986).

39. S. Leslie, et al., The fine-scale genetic structure of the British population. Nature 519, 309–314 (2015).

40. J. F. Edwards, B. P. Hindle, The transportation system of medieval England and Wales. J. Hist. Geogr. 17, 123–134 (1991).

41. K. D. Snell, English rural societies and geographical marital endogamy, 1700-1837. Econ. Hist. Rev. 55, 262–298 (2002).

42. B. W. Domingue, J. Fletcher, D. Conley, J. D. Boardman, Genetic and educational assortative mating among US adults. Proc. Natl. Acad. Sci. U. S. A. 111, 7996–8000 (2014).

43. N. Creanza, O. Kolodny, M. W. Feldman, Cultural evolutionary theory: How culture evolves and why it matters. Proc. Natl. Acad. Sci. U. S. A. 114, 7782–7789 (2017).

44. V. Labeyrie, M. Thomas, Z. K. Muthamia, C. Leclerc, Seed exchange networks, ethnicity, and sorghum diversity. Proc. Natl. Acad. Sci. U. S. A. 113, 98–103 (2016).

45. D. J. Lawson, G. Hellenthal, S. Myers, D. Falush, Inference of population structure using dense haplotype data. PLoS Genet. 8, e1002453 (2012).

