## Supplemental Tables and Figures for "Fine-scale cultural variation reinforces genetic structure in England"

PNAS

**This PDF file includes:**

Figures S1 to S6

Tables S1 to S3

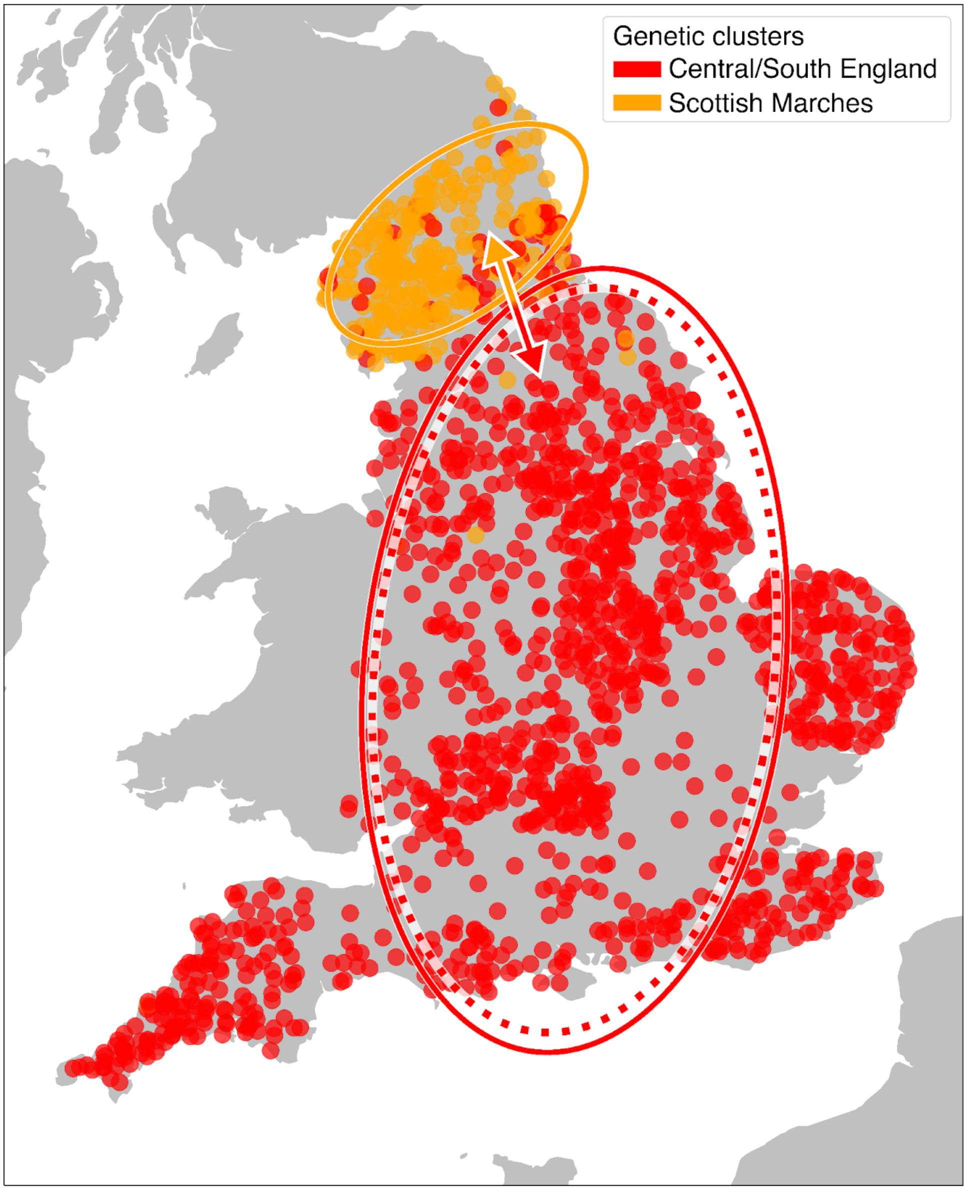


Fig. S1 **Relationship between rates of genetic and linguistic rates of change for 2 genetic clusters.** See Fig. 2, Tables S1 and S2. The gene-language relationship between the two clusters is significant. We draw a dotted ellipse in the cluster that represents much of the Midlands and South England, where there is a significant correlation between the spatial distributions of genes (of individuals in that cluster) and language (in the region where the cluster is found).

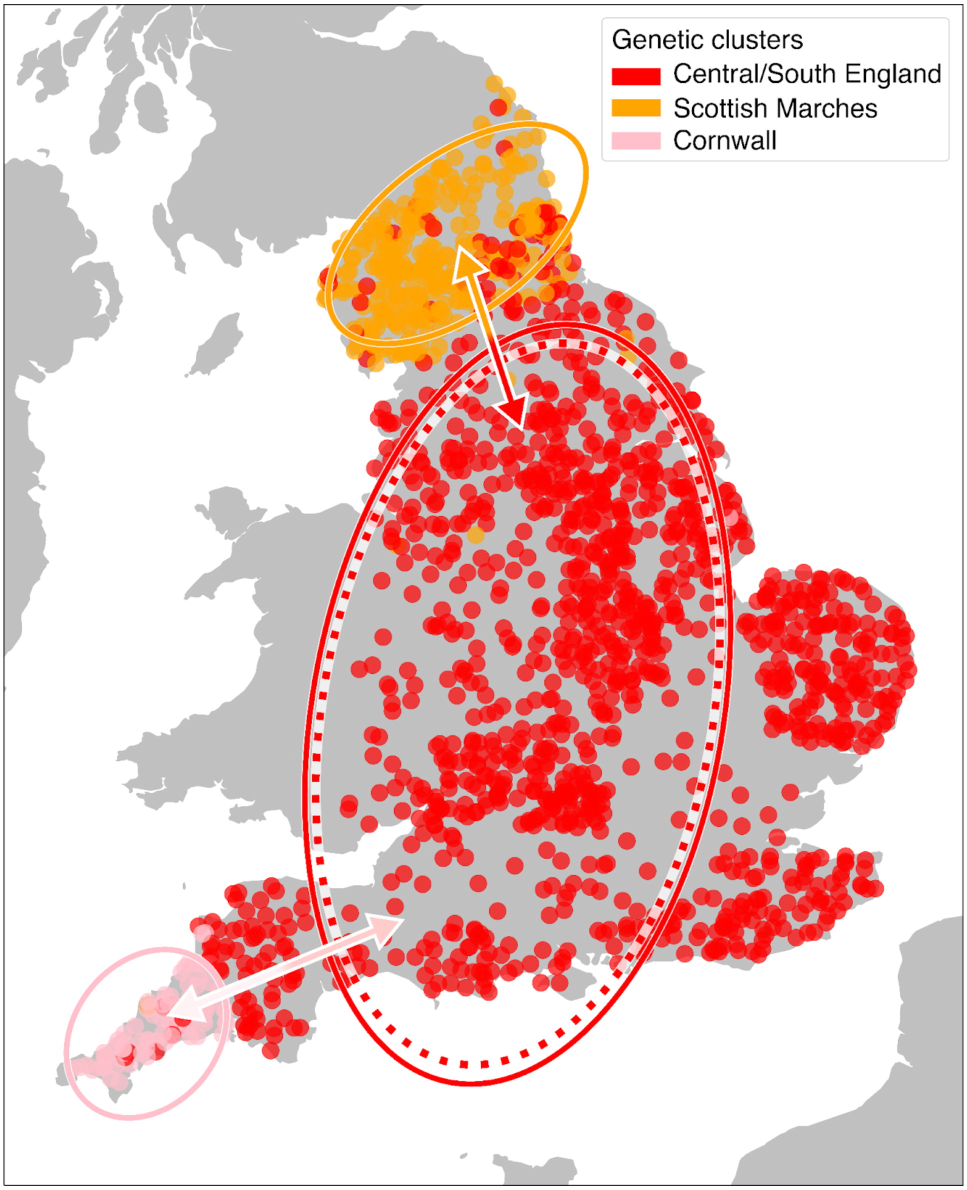


Fig. S2 **Relationship between rates of genetic and linguistic rates of change for 3 genetic clusters.** See Fig. 2, Tables S1 and S2. For between-cluster boundaries (double-sided arrows), a significant gene-language relationship is shown by a dark arrow and a non-significant relationship is shown by a light arrow. We draw a dotted ellipse in the cluster that represents much of the Midlands and South England to indicate that there is a significant correlation between the spatial distributions of genes (of individuals in that cluster) and language (in the region where the cluster is found).

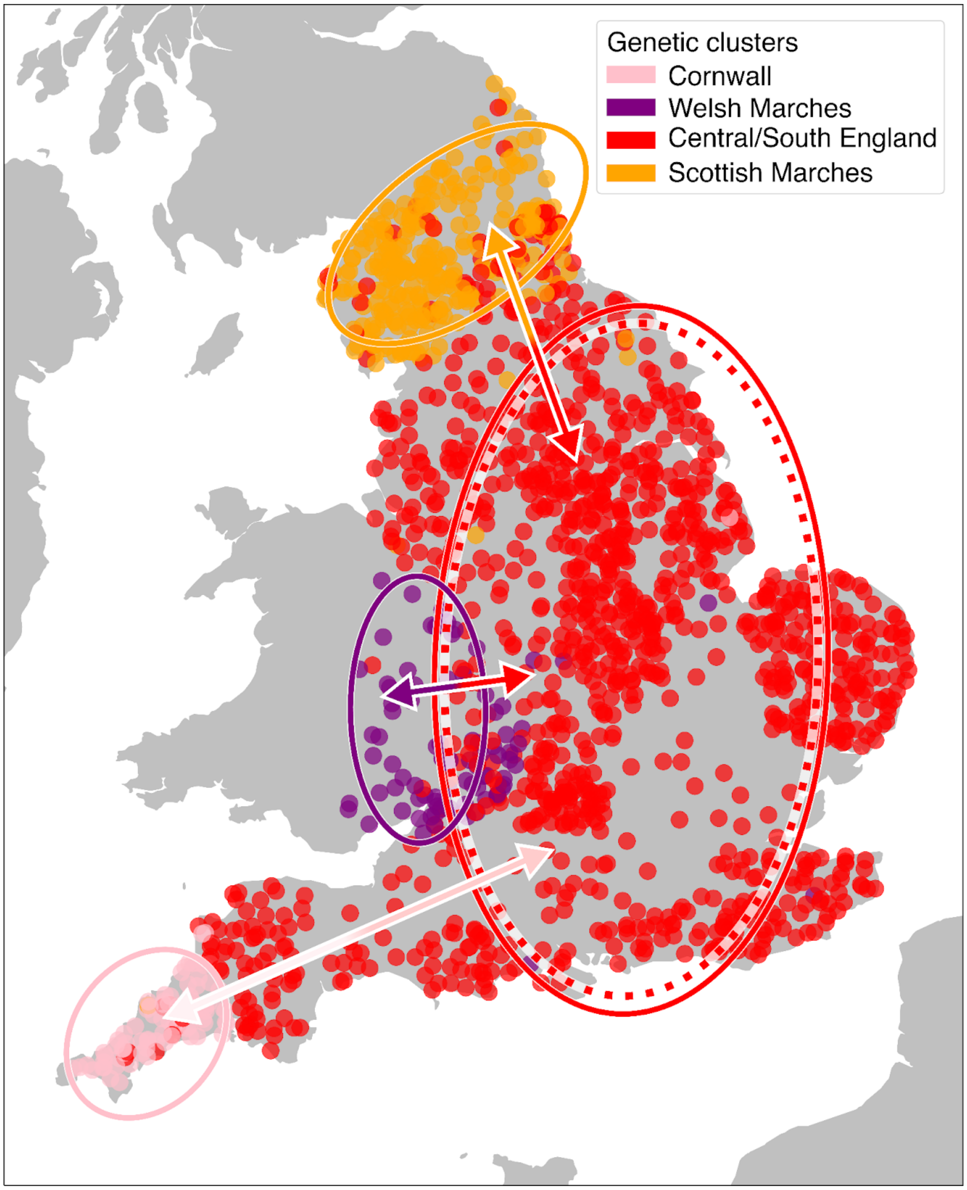


Fig. S3 **Relationship between rates of genetic and linguistic rates of change for 4 genetic clusters.** See Fig. 2, Tables S1 and S2. For between-cluster boundaries (double-sided arrows), a significant gene-language relationship is shown by a dark arrow and a non-significant relationship is shown by a light arrow. We draw a dotted ellipse in the cluster that represents much of the Midlands and South England to indicate that there is a significant correlation between the spatial distributions of genes (of individuals in that cluster) and language (in the region where the cluster is found).

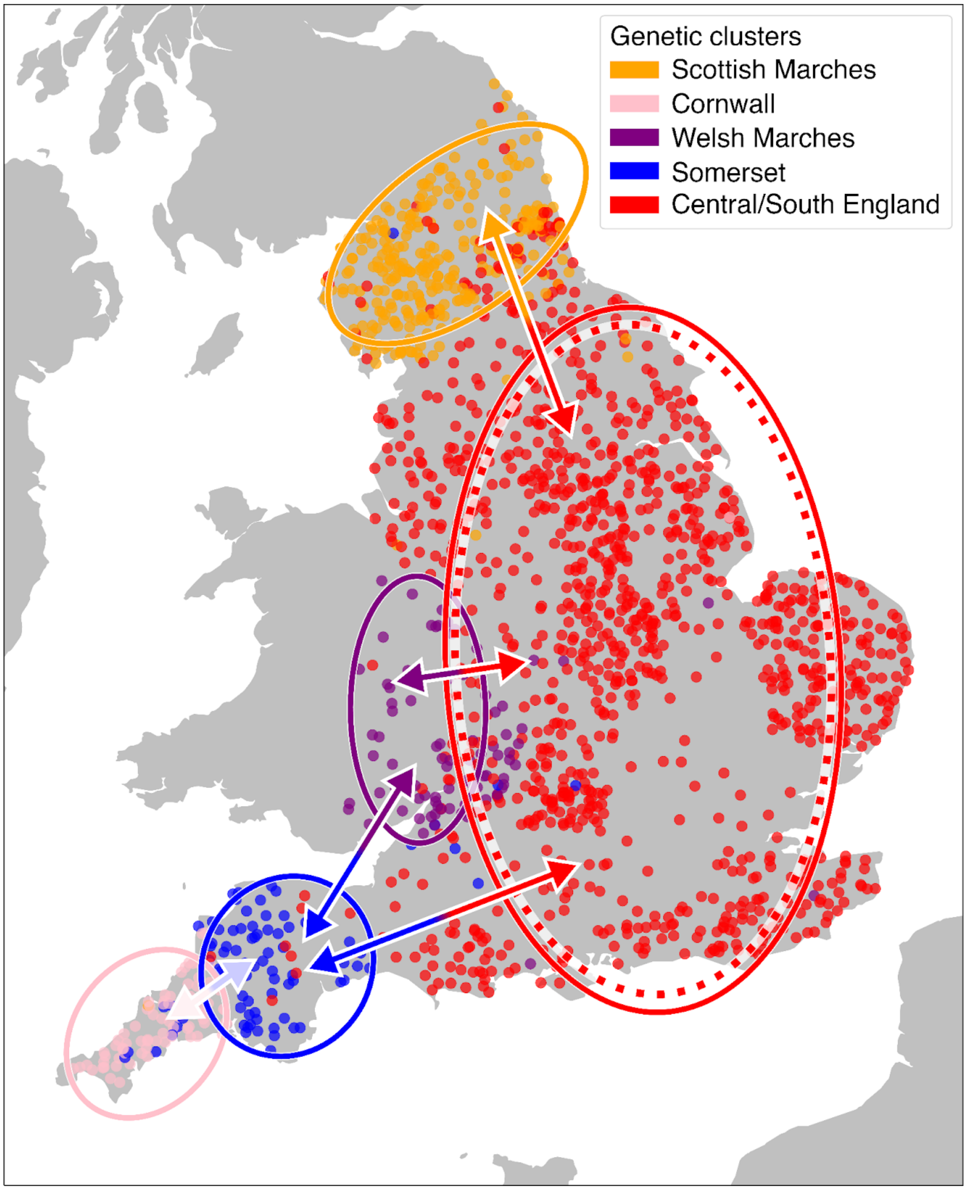


Fig. S4 **Relationship between rates of genetic and linguistic rates of change for 5 genetic clusters.** See Fig. 2, Tables S1 and S2. For between-cluster boundaries (double-sided arrows), a significant gene-language relationship is shown by a dark arrow and a non-significant relationship is shown by a light arrow. We draw a dotted ellipse in the cluster that represents much of the Midlands and South England to indicate that there is a significant correlation between the spatial distributions of genes (of individuals in that cluster) and language (in the region where the cluster is found).

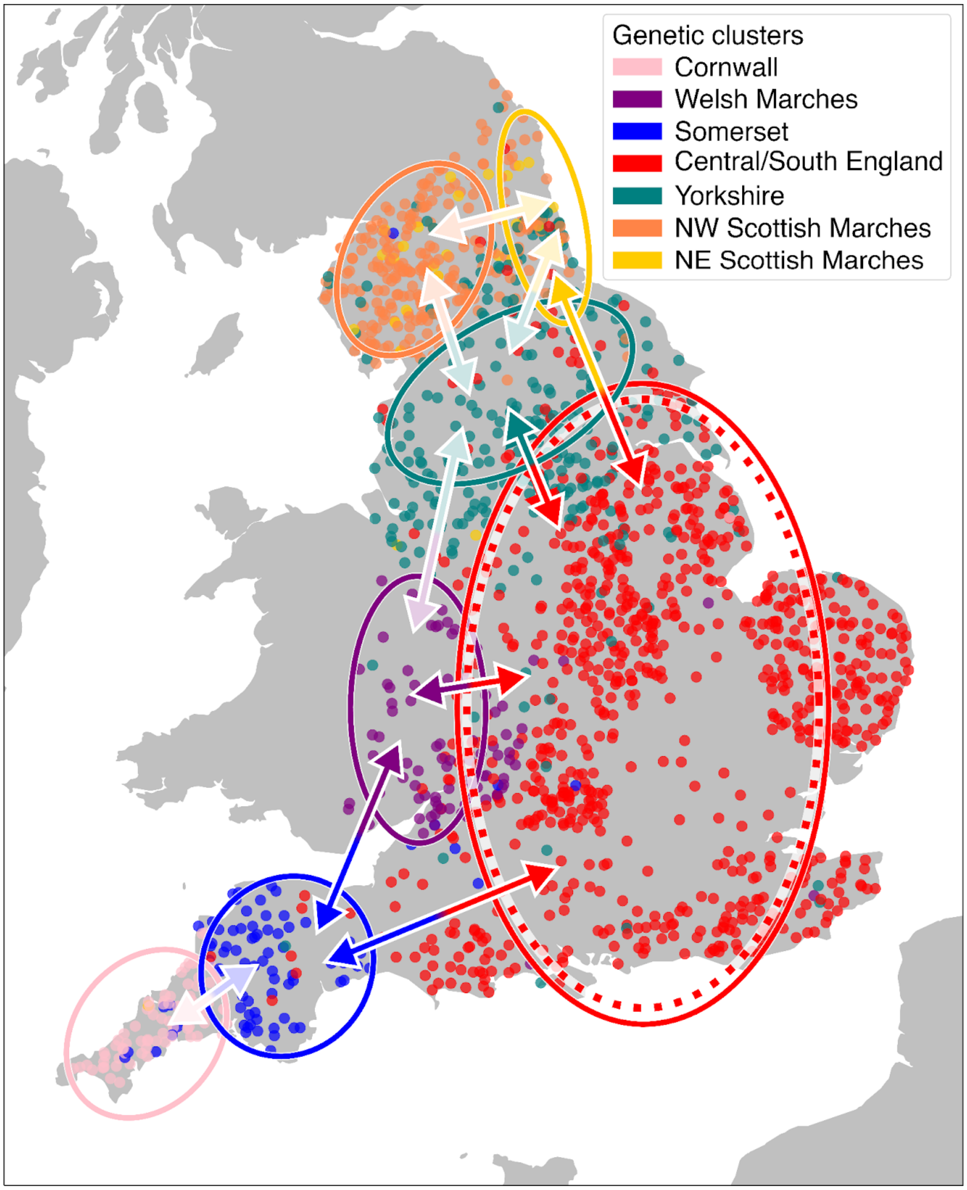


Fig. S5 **Relationship between rates of genetic and linguistic rates of change for 7 genetic clusters.** See Fig. 2, Tables S1 and S2. For between-cluster boundaries (double-sided arrows), a significant gene-language relationship is shown by a dark arrow and a non-significant relationship is shown by a light arrow. We draw a dotted ellipse in the cluster that represents much of the Midlands and South England to indicate that there is a significant correlation between the spatial distributions of genes (of individuals in that cluster) and language (in the region where the cluster is found).

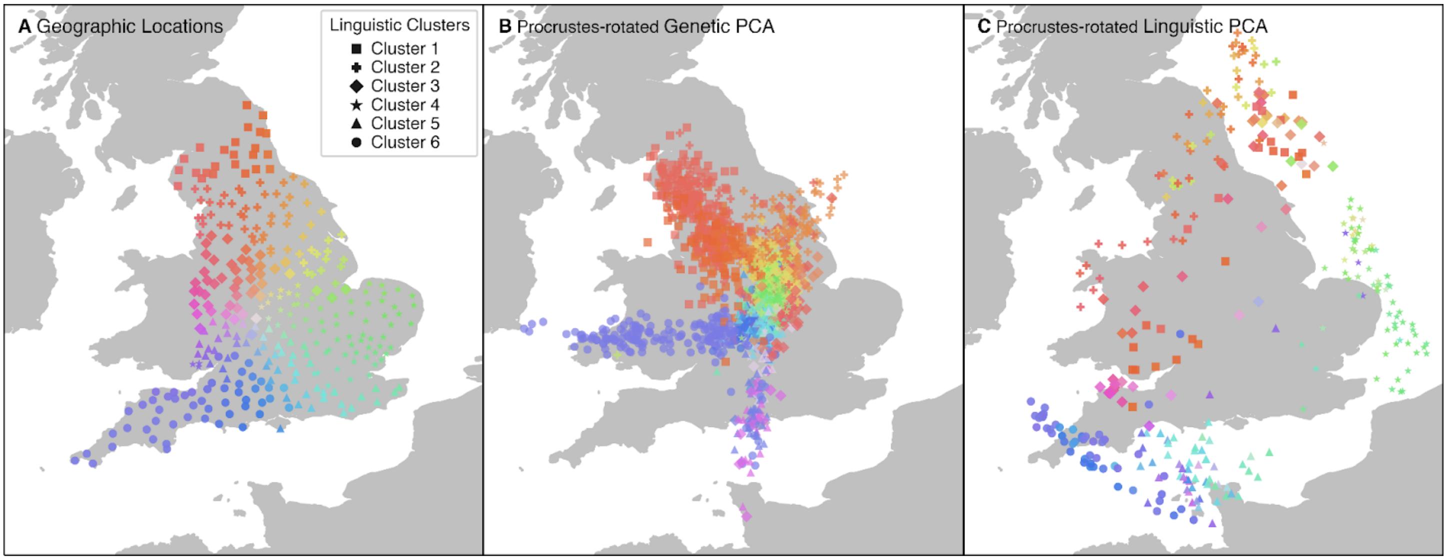


Fig. S6 **Procrustes transformations of genetic and linguistic variation on geography.** We use the same geographic coloring (A) to visualize genetic (B) and linguistic (C) variation after a Procrustes transformation of their first 3 principal components. See Fig. 3 and the main text for details. The Procrustes correlations are 0.683 for the genetics-geography analysis (B), 0.888 for language-geography (C), and 0.531 for language-genetics (not shown). All are significant using permutation tests (empirical $p < 10^{-6}$).

Table S1  **Association between the 55 linguistic feature categories and the words from SED used to construct them**.

| Feature | Words used |
| --- | --- |
| Fronted back half-open vowel ɔː (to æ/ɑ) | bald |
| Raised front open vowel a (to ɛ) | ladder |
| Front open vowel in aɪ diphthong (ɛɪ to aɪ) | fight |
| Diphthong ɑɪ fronted to aɪ | flies |
| Monophthongization of əʊ | coulter, counter, cow |
| Elision of b in bl̩ | thimble |
| Palatalized Old English k | thatcher, thatch, flitch, birch |
| Elision of d in nd/ld | bellyband, grindstone, bald, blind, hand, cold |
| Voicing of f in fs to vs | hoofs |
| Elision of l in lk | yolks |
| Elision of s (spz to psiz) | wasps |
| Elision of t in coda st | harvest, breast, waistcoat |
| Fortition of gl to dl | glove |
| Fortition of θ to f | thatcher, thatch |
| Fortition and stopping of θ to d | thatcher, thatch |
| Lowered ɛː to aː (hair vowel) | barefoot |
| Glottalization of t in tl̩ | kettle |
| Breaking of ɛː | chair, wear |
| Breaking of eɪ | spade, drain, gate, chain, knife |
| Monophthongization of eɪ | spade, hay, drain, gate, lay, chain |
| Voicing of initial f to v | knife, forks, fellies, foal, flitch, flight, ford, fox, fleas, flies, fern, useful, floor, fire, finger, foot |
| Glottalization of coda t | snout, quilt, suet, foot |
| Foot-Strut split | gums, duck |
| NG-coalescence | milkingstool, boiling, darning |
| H-loss (to f) | trough |

| | |
|---|---|
| H-dropping | halter, hammer, harvest, hay, hoof, hive, hollybush, head, hair, hear, hand, height, heat |
| H-insertion | arm, arse |
| Velarized l (to ɫ) | halter, saddle, bellyband, ladder, plough, coulter, wheel, stool, foal, colt, wool, slaughter, flitch, coal, lead, weasel, squirrel, lay, yolk, owl, lice, fleas, flies, hollybush, useful, quilt, oil, floor, shovel, gruel, kettle, shelf, needle, thimble, bald, flind, shoulder, nail, measles, cold, gloves |
| Elision of initial g | gate |
| Monophthongization of ɪə | hear |
| Elision of l | coulter, shovel, gruel, needle, cold |
| Lentition of medial tʃ | scratching |
| Raising and backing a | calf |
| Diphthongization of ɔː to ʊɔ | boar, ford, road, door, floor |
| Monophthongization of oʊ | road, cold |
| Unstressed ʊə | grindstone |
| Realization of ɔʊ as uː | ewe, coal |
| Splitting of oʊ diphthong | toad |
| Rhoticity | halter, girth, hammer, ladder, forks, coulter, cartgrease, turnips, harvest, thatcher, water, boar, butcher, slaughter, ford, bird, partridge, worms, birch, fern, pear,door, floor, chair, fire, cucumber, sugar, darning, hair, hear, shoulder, arm, finger, arse, barefoot, warts, thirsty |
| Voicing of s | saddle, sawdust, suck, sow, snout, useful, suet, sew, see, sweat |
| Lentition of ʃ | sugar |
| Fortition of ð to d | thimble, thresh |
| Toe-tow merger | meadow, mow |
| Lowering of mid-back mid-open vowel ʌ | drunk |
| Raising of unstressed ə | barrel, gloves |

| | |
|---|---|
| Metathesis of əʊ to ʊə | comb, nose, waistcoat |
| Elision of v | shovel |
| Elision of unstressed vowels | grindstone, naked |
| Whine-wine merger | wheel |
| Monophongization of oʊ | nose |
| Yod-insertion | cabbage, carrots, suet, kettle, calf |
| Yod-dropping | cucumber |
| Yod-coalescence | tune |

Table S2 **Genetic-linguistic covariation between clusters**. For each set of adjacent clusters at each number of hierarchical clusters (*K*), we measured whether their cluster boundary was found in regions with high linguistic rates of change using Mann-Whitney *U* tests. Bold values indicate those significant after Bonferroni correction.

| *K* | Cluster 1 | Cluster 2 | *p*-value | *K* | Cluster 1 | Cluster 2 | *p*-value |
|---|---|---|---|---|---|---|---|
| 2 | S/C. England | Scot. Marches | **$1\times10^{-5}$** | 6 | Yorkshire | Scot. Marches | $1.76\times10^{-2}$ |
| 3 | S/C. England | Scot. Marches | **$1\times10^{-5}$** | 6 & 7 | S/C. England | Scot. Marches | **$3\times10^{-5}$** |
| 3 | S/C. England | Cornwall | 0.293 | 6 & 7 | S/C. England | Yorkshire | **$8\times10^{-5}$** |
| 4 | S/C. England | Scot. Marches | **$<1\times10^{-5}$** | 6 & 7 | Yorkshire | Welsh Marches | 0.999 |
| 4 | S/C. England | Welsh Marches | **$<1\times10^{-5}$** | 6 & 7 | S/C. England | Welsh Marches | **$<1\times10^{-5}$** |
| 4 | S/C. England | Cornwall | 0.242 | 6 & 7 | S/C. England | Somerset | **$<1\times10^{-5}$** |
| 5 | S/C. England | Scot. Marches | **$<1\times10^{-5}$** | 6 & 7 | S/C. England | North East | **$2\times10^{-5}$** |
| 5 | S/C. England | Welsh Marches | **$<1\times10^{-5}$** | 7 | Yorkshire | North East | 0.021 |
| 5 | S/C. England | Somerset | **$1\times10^{-5}$** | 7 | Yorkshire | North West | $9.17\times10^{-3}$ |
| 5–7 | Somerset | Cornwall | 0.99 | 7 | North East | North West | 0.524 |
| 5–7 | Somerset | Welsh Marches | **$1.6\times10^{-4}$** | | | | |

Table S3 **Genetic-linguistic covariation within clusters**. For each cluster at each number of hierarchical clusters (*K*), we compared genetic and linguistic rates of change using linear regression. The only cluster that was significant for this analysis was the one encompassing South and Central England at all levels of hierarchical clustering (represented in red in Fig. 2). Bold values are those significant after Bonferroni correction. We also include the number of individuals in each cluster (*N*).

| *K* | Cluster (*N*) | *p*-value | *K* | Cluster (*N*) | *p*-value |
|---|---|---|---|---|---|
| 2 | S/C. England (1,372) | **$2.73\times10^{-8}$** | 5 | S/C. England (1,122) | **$1.21\times10^{-12}$** |
| 2 to 6 | Scottish Marches (295) | 1 | 5 to 7 | Somerset (82) | 0.977 |
| 3 | S/C. England (1,290) | **$1.44\times10^{-12}$** | 6 to 7 | S/C. England (836) | **$3.14\times10^{-11}$** |
| 3 to 7 | Cornwall (82) | NA (single county) | 6 to 7 | Yorkshire/NW England (286) | $5.91\times10^{-2}$ |
| 4 | S/C. England (1,204) | **$5.56\times10^{-12}$** | 7 | North East England (32) | 1 |
| 4 to 7 | Welsh Marches (86) | 3.66e-2 | 7 | North West England (263) | 1 |
